# Genetic diversity, agroecosystems, and climate suitability of cacao in Panama: historical legacies and conservation priorities

**DOI:** 10.64898/2026.11.12.764092

**Authors:** Hilda E. Castillo, David Martínez Joven, Luis C. Mejía

## Abstract

*Theobroma cacao* is a strategic resource for socioeconomic development and ancestral knowledge preservation in Panama. We integrated SNP genotyping, historical analysis, and climate suitability modeling to characterize the genetic resources of Panamanian cacao and inform future conservation priorities.
We genotyped 153 cacao accessions from eight provinces using 96 SNP markers, assessed genetic diversity and ancestry composition, evaluated associations between ancestry and collection habitat (farms, forests, and home gardens), and projected current and future (2050, SSP5-8.5) climatic suitability using species distribution models.
Genetic diversity was moderate (He = 0.384) with high admixture (54.7%); Amelonado (46.3%) and Criollo (29.1%) predominated. Criollo ancestry was strongly associated with home gardens and, to a lesser extent, forests, relative to farms (Firth-adjusted OR = 76.8, p < 0.001, and OR = 20.7, p = 0.013), whereas Amelonado predominated in commercial farms, reflecting pre-Columbian dispersal, colonial persistence, commercial introductions, and modern breeding. Climatically suitable area is projected to increase 11.9% by 2050 (8,780 to 9,825 km²), with gains in Chiriquí, Veraguas, and northwestern Darién, and losses along the Pacific coast of Veraguas and southwestern Los Santos.
Human-managed agroecosystems, particularly home gardens, play a disproportionate role in maintaining Criollo-associated ancestry in Panama, including Criollo-associated genotypes not currently held in international reference collections. We recommend a national germplasm bank and the integration of genetic and climate information into conservation and adaptation strategies for Panamanian cacao.

**Societal Impact Statement:** Cacao has supported rural livelihoods, cultural traditions, and Indigenous knowledge systems in Panama for centuries. However, climate change and the erosion of traditional agroecosystems threaten both cacao diversity and the communities that maintain it. Our study shows that home gardens and forest remnants harbor disproportionate levels of Criollo-associated diversity, whereas farms maintain mainly Amelonado and admixed germplasm, while also identifying regions that may remain climatically suitable for cultivation in the future. Protecting these genetic resources, together with the cultural practices that sustain them, can strengthen climate-resilient cacao production, support rural economies, and safeguard Panama’s biocultural heritage for future generations.

## 1 Introduction

Cacao (*Theobroma cacao* L.), the source of chocolate, is a globally important crop whose sustainability depends on the conservation of genetic diversity and its capacity to adapt to environmental change. Across tropical regions, cacao production supports millions of smallholder farmers and sustains complex agroecosystems shaped by longstanding interactions between ecological processes and human management (Bartley, 2005; Coe & Coe, 1996). However, emerging diseases and the progressive erosion of genetic resources pose significant threats to the future of cacao cultivation, particularly in regions where diversity remains poorly characterized.

Genetic diversity underpins crop resilience and breeding for disease resistance and productivity (Ceccarelli et al., 2022; Nieves-Orduña et al., 2023). In cacao, this diversity originated in the Upper Amazon Basin, where the species exhibits its highest genetic variation and where domestication processes began more than 5,000 years ago (Bartley, 2005; Zarrillo et al., 2018). From this center of origin, cacao was dispersed through complex networks of human migration and exchange, eventually reaching Mesoamerica where it became deeply embedded in cultural, economic, and ritual systems (Coe & Coe, 1996; Henderson et al., 2007; Powis et al., 2002). Recent archaeogenomic evidence suggests that this dispersal involved multiple genetic lineages, reflecting dynamic processes of selection, movement, and cultivation across landscapes (Lanaud et al., 2024).

Cacao was historically classified into two major genetic groups: Criollo, associated with Mesoamerica and valued for fine flavor but disease-susceptible, and Forastero, more prevalent in South America and Africa and valued for resilience and yield (Motamayor et al., 2008). Hybridization of these groups produced Trinitario cacao, now central to modern cacao production and breeding (Lanaud et al., 2024).

Using SSR markers, Motamayor et al. (2008) classified cacao germplasm into ten major genetic groups, including Amelonado, Contamana, Criollo, Curaray, Guiana, Iquitos, Marañón, Nacional, Nanay, and Purús. Subsequent studies have identified additional regional diversity in Ecuador and Colombia (Argout et al., 2023), Bolivia (Zhang et al., 2012), Peru (Arevalo-Gardini et al., 2019), highlighting the dynamic and geographically structured nature of cacao genetic diversity.

Panama occupies a unique position within the historical dispersal of cacao, acting as a bridge between South and Central America. Archaeobotanical and archaeogenomic evidence confirms the pre-Columbian use of cacao and its persistence through colonial and post-colonial periods despite major disruptions in land use and population dynamics (Dickau, 2010; Lanaud et al., 2024; Castillero Calvo, 2019; Cooke & Sánchez, 2003). These long-term interactions between people and landscapes suggest that cacao diversity in Panama reflects both biological processes and centuries of cultural practices and agroecological management.

Traditional agroecosystems, including home gardens (small multispecies plots near homesteads) and smallholder farms (small-scale, family-managed plots), are increasingly recognized as important reservoirs of crop genetic diversity, where locally adapted lineages are maintained through low-intensity management and cultural selection (Galluzzi et al., 2010). In cacao, these systems may harbor unique genetic resources that are absent from formal germplasm collections, particularly in countries where national conservation strategies are limited. In Panama, the absence of a centralized cacao germplasm bank increases the risk of genetic erosion and constrains the use of local diversity for breeding and adaptation. Understanding how different agroecosystems contribute to the maintenance of genetic diversity is therefore critical for both conservation and sustainable production.

At the same time, climate change is expected to alter the distribution of suitable environments for cacao cultivation, potentially reshaping production landscapes and exacerbating existing vulnerabilities (Bunn et al., 2019; Ceccarelli et al., 2022). Species distribution models provide valuable insights into these shifts, but are rarely integrated with genetic and historical data, limiting their usefulness for identifying which genetic resources may be most relevant under future conditions (Sillero et al., 2021). Bridging this gap is essential for developing strategies that align conservation, production, and climate adaptation. A recent study characterized cacao genetic structure in Panama’s principal cacao-producing region, Bocas del Toro, and in Herrera, with sampling restricted to farms (Ferro-Lozano et al., 2026). Our study differs in both geographic coverage and the diversity of management systems examined, encompassing forests and home gardens in addition to farms across a broader set of provinces, while prioritizing naturalized trees without a documented history of recent introduction.

Here, we investigate how historically shaped agroecosystems in Panama contribute to the maintenance of cacao genetic diversity and assess their potential relevance under future climate scenarios. Specifically, we (i) characterize the genetic diversity and population structure of Panamanian cacao using SNP markers, (ii) evaluate the association between genetic ancestry and different management systems (farms, forests, and home gardens), (iii) contextualize these patterns within historical processes of dispersal and cultivation, and (iv) model current and future climatic suitability to identify regions of persistence and change.

## 2 Materials and methods

### 2.1 Sample Collection and DNA Extraction

Cacao samples were collected from naturalized seedling trees found in farms, private gardens, and forests throughout Panama. Trees were located through collaboration with local communities, natural reserves, and Panamanian institutions. Sampling was conducted with the consent of local farmers and landowners. All necessary permits for genetic resource collection and export were obtained from the relevant authorities in Panama.

A targeted sampling approach was implemented, prioritizing cacao trees without a known history of modern introduction. Passport information, phenotypic traits (pod shape, seed color, tree health), and environmental data were recorded for each tree using a structured questionnaire.

Total genomic DNA was extracted using DNeasy® Plant Mini kit (Qiagen Inc., Valencia, CA, USA) following the specified kit instruction.

### 2.2 SNP Markers and Genotyping

The single nucleotide polymorphism (SNP) markers used in this study were selected from expressed sequence tags (ESTs) generated from cDNA libraries representing diverse cacao plant tissues. A panel of 96 SNP loci, previously characterized was employed for genotyping (Lindo et al., 2018; Zhang et al., 2012).

Genotyping was conducted using the Fluidigm 96.96 Dynamic Array™ system (Fluidigm, San Francisco, CA), which allows high-throughput analysis of 96 samples against 96 SNP assays in parallel, yielding 9,216 genotype calls per run. To ensure robust amplification, a specific target amplification (STA) step was incorporated to enrich template DNA prior to multiplexed genotyping (Arevalo-Gardini et al., 2019; Lindo et al., 2018; B. Wang et al., 2020; J. Wang et al., 2009). The STA reaction was performed according to the manufacturer’s protocol, using a master mix consisting of 2.5 μL TaqMan® PreAmp Master Mix (2X) (Life Technologies, Carlsbad, CA).

### 2.3 Data Processing and Genetic Analyses

Raw genotyping data were compiled and filtered in Microsoft Excel®, excluding samples with >10% missing calls from subsequent analyses. Duplicate accessions were identified through pairwise multilocus comparison of SNP profiles using GenAlEx 6.5 (Peakall & Smouse, 2012), retaining only one representative per genetically identical group. For comparative analyses, we included a reference set of 219 cacao accessions with previously published SNP profiles (Arevalo-Gardini et al., 2019; Cosme et al., 2016; Lindo et al., 2018).

Genetic diversity parameters, including Shannon’s information index (I), observed heterozygosity (Ho), expected heterozygosity (He), minor allele frequency (MAF), fixation index (F), and inbreeding coefficient (FIS), were calculated using GenAlEx 6.5. Population structure was assessed using principal coordinate analysis (PCoA) based on SNP-derived genetic distances.

After converting SNP genotypes to a *genind* object using *adegenet*, pairwise genetic dissimilarities among accessions were calculated using the diss.dist function from the *poppr* package (Kamvar et al., 2014). A UPGMA dendrogram was constructed from the resulting dissimilarity matrix using *phangorn* (Schliep, 2011). Final visualization and annotation were performed with *ggtree* (Yu et al., 2017) using a circular layout. All analyses were conducted in R v4.3.2 (R Core Team, 2023).

Population structure was further investigated using a Bayesian clustering approach implemented in STRUCTURE v2.3.4 (Pritchard et al., 2000). The analysis incorporated 10 reference cacao populations representing major genetic groups of *T. cacao* (Motamayor et al., 2008). To mitigate potential bias from uneven sample sizes, each reference group was standardized to 200 individuals using simulated genotypes generated in ONCOR (Kalinowski et al., 2007). The admixture model was applied with K=10, and ten independent runs were performed, each consisting of 200,000 Markov chain Monte Carlo (MCMC) iterations following a 100,000-iteration burn-in period. The run with the highest log probability (Ln Pr(X|K)) was selected for interpretation.

Finally, we classified cacao samples into three genotypic categories using a Q-value threshold of 0.75: Criollo (Q ≥ 0.75), Amelonado (Q ≥ 0.75), and admixed (Q < 0.75). The purpose of this classification was to distinguish accessions with predominantly single-group ancestry from admixed individuals and to evaluate how ancestry composition varied across habitats and geographic gradients.

Associations between dominant ancestry and collection habitat (farm, forest, or home garden) were first explored using multinomial logistic regression with elastic net regularization (α = 0.5), implemented in the glmnet package (Friedman et al., 2010). Dominant ancestry was modeled as a function of habitat and mean-centered longitude with Farm set as the reference level for habitat, and the regularization parameter (λ) was selected via 10-fold cross-validation with a fixed random seed for reproducibility. Because glmnet’s multinomial family estimates coefficients jointly for all classes rather than relative to a single class fixed at zero, exponentiated coefficients from this model do not constitute valid class-versus-reference odds ratios and are not reported as such; this model was instead used to generate predicted ancestry probabilities across the observed habitat and longitudinal gradient (Figure S2).

Inspection of the habitat × ancestry contingency table showed that Criollo ancestry was entirely absent among farm accessions (0 of 64), producing quasi-complete separation (Albert & Anderson, 1984) in the corresponding binary comparison. Firth penalized logistic regression (Firth, 1993), implemented in the logistf package (Heinze & Schemper, 2002), was therefore used as the primary inferential model for the association between Criollo ancestry (versus all other groups) and habitat. To match the covariate structure of the descriptive multinomial model and account for the underlying longitudinal ancestry gradient, this model included both habitat and mean-centered longitude as predictors, obtaining bias-reduced coefficients, odds ratios, and profile-likelihood 95% confidence intervals valid under sparse or separated data. As a sensitivity check, we compared the habitat effect estimated with and without adjustment for longitude to evaluate the extent to which the unadjusted association reflected geographic confounding rather than habitat type per se. As a descriptive complement, the observed proportion of Criollo ancestry within each habitat was calculated together with exact Clopper–Pearson 95% confidence intervals (Clopper & Pearson, 1934).

### 2.4 Historical events in Panamá

To better understand the genetic diversity of cacao in Panama, we conducted a narrative historical synthesis of records identifying key events that influenced cacao cultivation. Research sources included scholarly literature, colonial trade records, botanical surveys, and agricultural reports.

Digital repositories such as Google Scholar, and national archives were consulted to track the introduction and dispersal of cacao.

### 2.5 Occurrence data compilation and curation

We assembled a database of 274 georeferenced presence records for *T. cacao* by integrating three complementary sources: publicly available herbarium and observation data from GBIF accessed through the rgbif R package; targeted field surveys conducted in 2022 and 2024; and peer-reviewed literature records. All records were projected to the WGS 84 datum, assigned unique identifiers, and merged into a single spatial layer. Duplicate coordinates and records lacking complete spatial metadata were removed, retaining one observation per locality.

To minimize geolocation errors, we applied a multi-step quality control workflow. Coordinate outliers were flagged with CoordinateCleaner, which screens for points located on country centroids, biodiversity institutions, or ocean pixels. Administrative information was standardized with the countrycode utility, and geometric checks such as distance to coast and topology tests were run with rgeos. Records that failed these filters were inspected manually and retained only when locality notes could be verified.

Spatial sampling bias was reduced by applying a 1-km rarefaction filter using thin_records() in EcoNicheS, which wraps the distance-based spThin algorithm. Finally, 400 pseudo-absence points were generated using a 1:2 presence-to-absence ratio (Barbet-Massin et al., 2012).

### 2.6 Bioclimatic predictors

We assembled the 19 BIOCLIM variables at 30 arc-second resolutions from WorldClim v2.1 (Fick & Hijmans, 2017). Contemporary climatology represented the 1970–2000 baseline, and mid-century projections represented 2041–2060 under the high-emissions SSP5-8.5 scenario. Mid-century projections were selected because they align with climate-smart agriculture planning horizons for food security (Lipper et al., 2014). For the future projection, we retained the 13 general circulation models that provided the complete BIOCLIM suite for this scenario. Data processing was carried out in R using geodata (Hijmans et al., 2021), terra (Hijmans et al., 2020), countrycode (Arel-Bundock et al., 2018), ggplot2 (Wickham et al., 2025), reshape2 (Wickham, 2007),and usdm (Naimi, 2012). To avoid over-parameterisation, we first calculated pairwise Pearson correlations and, within each highly correlated pair, retained the predictor with the strongest ecological relevance for the target species. We then computed variance-inflation factors in usdm and removed layers with values greater than 10 to reduce multicollinearity and avoid inflated regression standard errors. The final set of retained predictors is provided in Table S7, and model performance statistics are reported in Table S8. These variables were used to calibrate the climatic suitability models.

### 2.7 Climatic suitability modeling

Climatic suitability for the baseline climate and for 2050 was estimated with the R package EcoNicheS, which integrates data acquisition, ecological-niche modeling, species-distribution modeling and landscape-connectivity analysis into a single workflow (Sunny et al., 2025). Presence records from GBIF and contemporary climate layers were retrieved through this package. Model calibration, evaluation and ensemble forecasting were carried out with the BIOMOD2 framework (Thuiller et al., 2025). Two machine-learning algorithms Random Forest and MaxEnt were selected because they capture complex, non-linear relationships between predictors and occurrences (Osorto Nuñez, 2022; Zhao et al., 2022). Occurrence data were split at random into training (80 %) and validation (20 %) subsets. Predictive performance was assessed with the area under the receiver-operating- characteristic curve (ROC), Cohen’s Kappa and the True Skill Statistic (TSS) (Sillero et al., 2021). Both Kappa and TSS range from −1 to 1: values close to 1 indicate high agreement between predictions and observations, whereas values near 0 or below denotes performance no better than random. We ran 10 independent replicates and applied a ROC threshold of 0.4 for binary conversion (Huang et al., 2023; Sunny et al., 2025).

Model robustness was further examined with Partial ROC (Peterson et al., 2008), which contrasts model predictions against null expectations while penalizing over-prediction. We accepted a 5% omission threshold for randomly withheld points in each subsample and executed 500 iterations to compute the partial area under the curve. The ratio of model AUC to the null expectation must exceed 1 to indicate statistical significance. We report the mean partial AUC across all iterations together with the corresponding ratios, where a value of 1 denotes random performance. These diagnostics support model performance, but they should be interpreted cautiously because high discrimination metrics can also reflect spatial sorting, sampling bias, or overly easy pseudo-absence contrasts.

### 2.8 Climatic suitability change

We quantified the change in climatic suitability expected for 2050 by directly comparing the continuous suitability surfaces generated for the baseline period and the future scenario. The future suitability raster was reprojected and resampled so that its resolution, spatial extent, and coordinate reference system matched those of the baseline map. We then subtracted baseline values from future values at the pixel level to produce a raster of absolute suitability change. A relative threshold equal to a fixed percentage of the overall suitability range was applied to classify each cell as loss, gain, or retention.

## 3 Results

### 3.1 Historical context shaping cacao diversity in Panama

The history of cacao cultivation in Panama reflects a long process of human-mediated dispersal, cultivation, exchange, and agricultural transformation. Archaeological and archaeogenomic evidence indicates that cacao was present and used in Panama well before European contact. Residues of *Theobroma cacao* DNA and methylxanthines recovered from ceramic vessels dated between 1500 BCE and 600 CE confirm the early use of cacao in the region and Panama (Lanaud et al., 2024). Additional evidence from Los Santanas and Río Bermejito in Coclé and Veraguas provinces of Panama further supports the presence of cacao beyond its native lowland range, suggesting transport, cultivation, or exchange by Indigenous communities (Dickau, 2010). Archaeogenomic analyses further revealed the presence of multiple cacao genetic lineages, including Amelonado, Nacional, and Purús related ancestries, suggesting that cacao diversity in Panama was already complex during pre-Columbian times (Lanaud et al., 2024). Together, these findings indicate that cacao formed part of regional exchange networks and cultural practices long before European colonization, creating opportunities for the movement and mixing of genetic lineages across western Panama (Figure 1).

**Figure 1.**
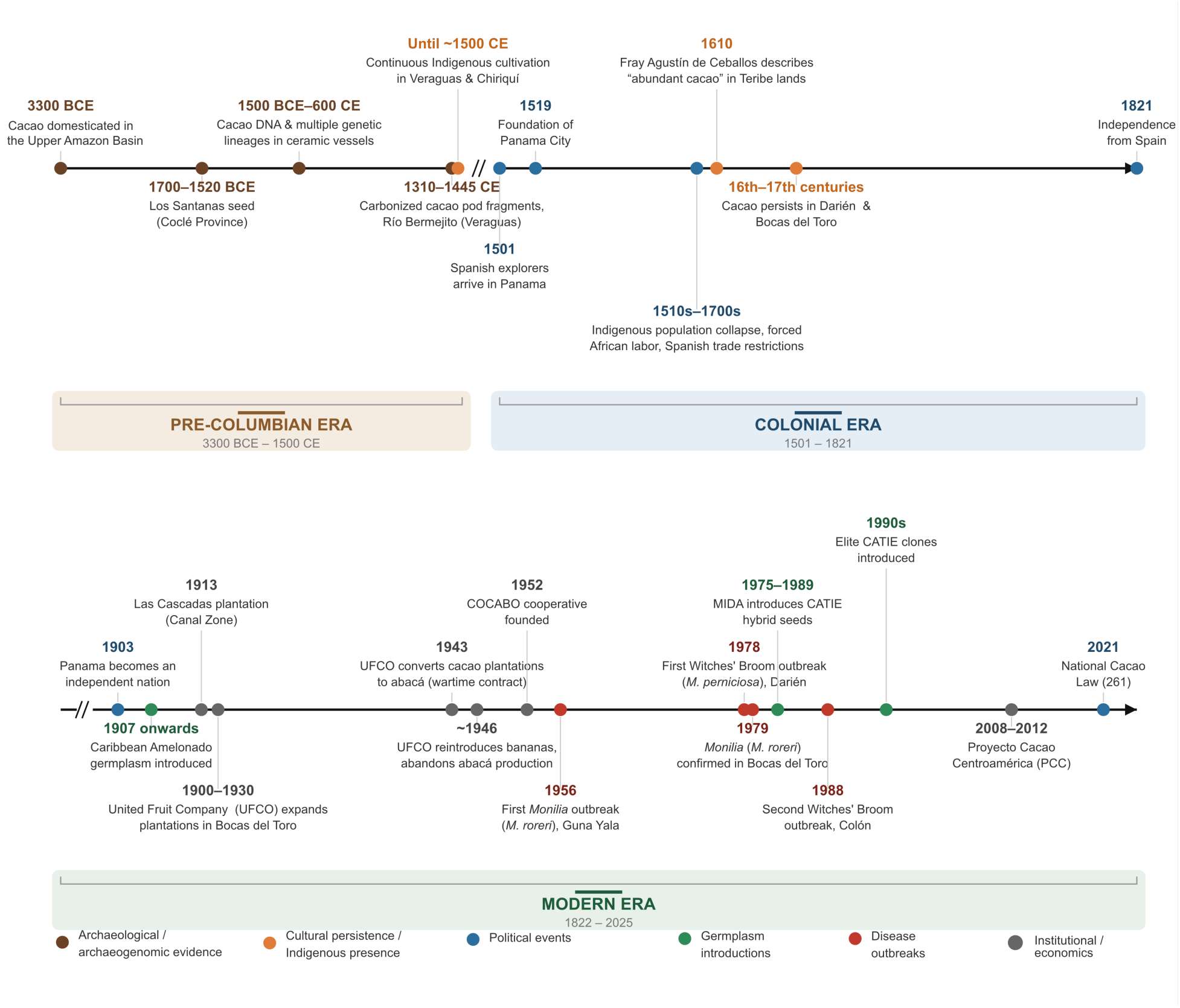
Historical timeline of cacao cultivation, germplasm introductions, disease outbreaks, and development milestones in Panama (3300 BCE–2025), shown across two rows: Pre-Columbian/Colonial eras above, Modern Era below. Colors indicate event category (see legend). Diagonal breaks (//) mark compressed axis segments. Additional sources: Bourgois (1994), Pearson & Holden (1945), COCABO (2025), CATIE (2018), Phillips-Mora (2003), Phillips-Mora et al. (2007), Gutiérrez (2020), Gobierno de Panamá, (2022); see Section 3.1.

Historical records suggest that cacao cultivation persisted throughout the colonial period in regions such as Veraguas, Chiriquí, Darién, and Bocas del Toro. Indigenous groups maintained cacao within agroecological production systems despite major demographic and social disruptions associated with Spanish colonization, including population decline, forced labor, and land-use change (Castillero Calvo, 2017; Fernández de Oviedo, 1851; Reverte Coma, 1995). In semi-autonomous regions such as Darién and Bocas del Toro, Indigenous and Afro-descendant communities likely contributed to the persistence of cacao cultivation despite broader colonial disruptions. The continued presence of cacao in these regions highlights the long-term role of local communities in maintaining cacao trees across changing political and ecological contexts.

From the late nineteenth century onward, cacao cultivation in Panama became increasingly influenced by commercial agriculture and germplasm introductions. Large-scale plantations were established in the Canal Zone and Bocas del Toro, where Forastero-derived Amelonado materials introduced from Trinidad and Tobago and other Caribbean sources became widely planted (Bennett & Taylor, 1912; Mata-Quirós et al., 2018; The Canal Record, 1914) (Figure 2D–E). During the twentieth century, successive outbreaks of frosty pod rot (*Moniliophthora roreri*) and witches’ broom (*Moniliophthora perniciosa*) triggered extensive rehabilitation programs and the introduction of improved germplasm through breeding initiatives involving CATIE and national agricultural programs (Cerda & Orozco, 2009; Phillips-Mora et al., 2007). These efforts introduced diverse genetic materials aimed at increasing productivity and disease resistance across cacao-growing regions.

**Figure 2.**
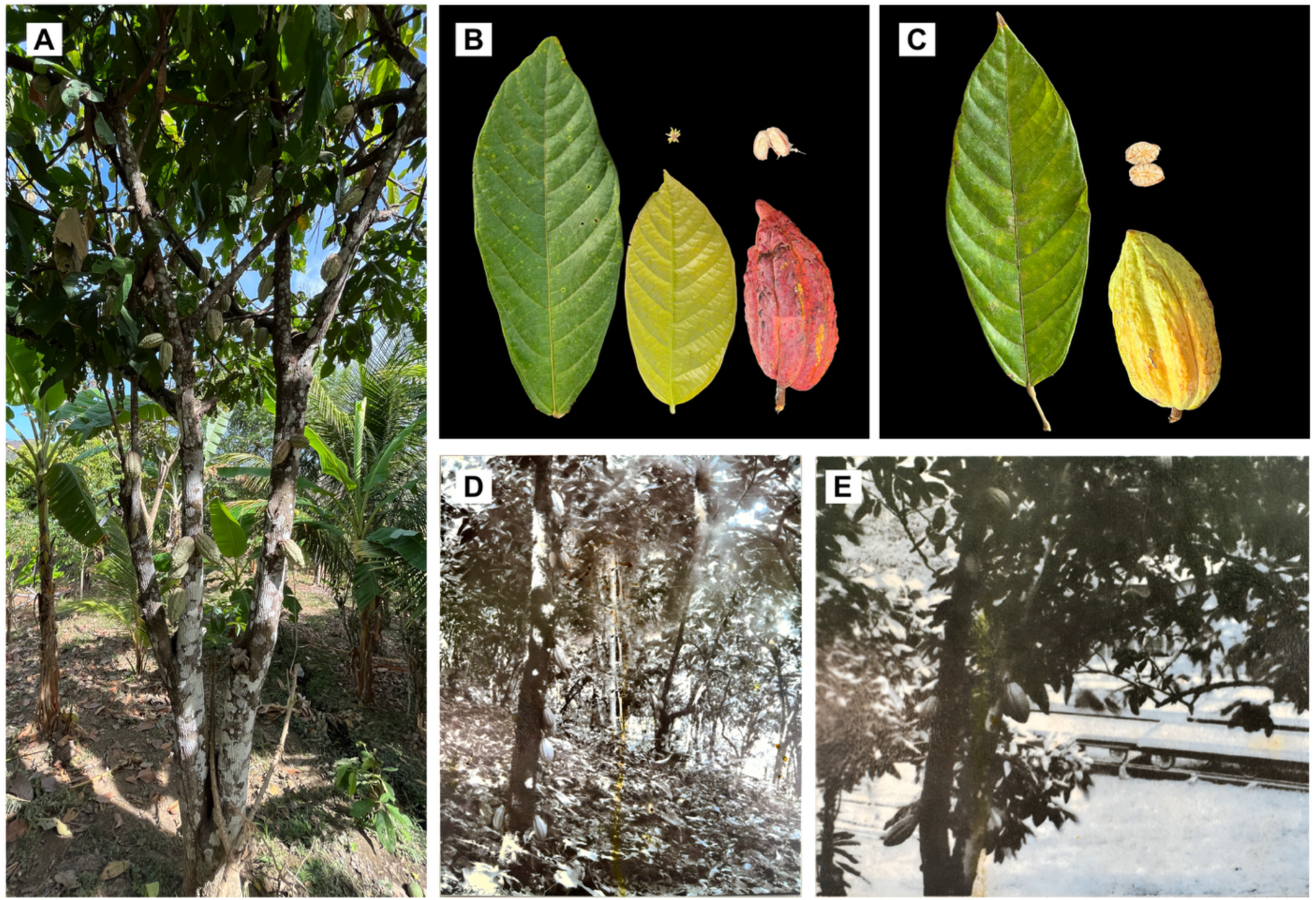
Contemporary and historical records of *Theobroma cacao* in Panama. (A) Criollo tree bearing developing pods, maintained in a home garden. (B) Cacao material from a Criollo tree in a forested area of Darién Province, including mature leaf, young leaf, flower, ripe red pod, and white cotyledon seeds. (C) Cacao material from a Criollo-type tree in Veraguas Province, including mature leaf, ripe yellow pod, and white cotyledon seeds. (D) Early twentieth-century photograph of cacao cultivation at Las Cascadas plantation during the Panama Canal construction period. (E) Archival photograph of cacao bean drying trays at the same plantation. Photo credits: (A–C) H.E. Castillo; (D–E) reproduced from interpretive signage at Plantation Road, Panama, displayed by the Autoridad Nacional del Ambiente.

Today, cacao remains economically and culturally important in Panama, particularly in Bocas del Toro and Indigenous territories, where traditional cultivation practices coexist with improved varieties. These overlapping historical processes of ancient dispersal, colonial persistence, commercial introductions, and modern breeding likely explain the complex ancestry and admixture patterns observed in contemporary Panamanian cacao populations. Major events related to cacao cultivation, exchange, disease outbreaks, and germplasm introductions are summarized in Figure 1.

### 3.2 SNP Genotyping, Duplicate Identification and Descriptive Statistics

A total of 153 cacao accessions were sampled from eight provinces across Panama, with the highest numbers collected from Bocas del Toro (n = 51) and Herrera (n = 51), followed by Colón (n = 12), Panamá (n = 11), Chiriquí (n = 11), Darién (n = 10), Coclé (n = 4), and Veraguas (n = 3) (Figure S1).To minimize the influence of recent introductions, we prioritized trees from older sections of farms, abandoned plantations, private gardens, and adjacent forested areas. Genotyping was conducted using 96 SNP markers, all of which passed quality thresholds (>90% call rate) and were retained for multilocus comparisons. One sample was excluded due to excessive missing data, resulting in 152 accessions for downstream analyses. Pairwise multilocus matching identified 32 accessions (20.9%) as genetic duplicates, grouped into eight synonymous clusters (Table S1). These groups consisted of individuals sharing identical SNP profiles across all 96 loci. From each cluster, a single representative was retained for downstream analyses, resulting in a filtered dataset of 128 unique genotypes.

The highest proportion of duplicate genotypes was found in Herrera, where approximately 47.0% of the sampled trees (24 of 51) were genetically redundant. In Darién, 4 of 10 accessions were duplicates, while Chiriquí and Bocas del Toro each contained two duplicated genotypes. No duplicates were identified in Panamá, Coclé, Colón, or Veraguas, although these provinces had smaller sample sizes.

Using the Panamanian cacao samples, we calculated summary statistics for 96 SNP markers (Table S2). Shannon’s information index (I) averaged 0.57, reflecting both allele richness and evenness, and ranged from 0.096 (locus TcSNP277) to 0.693 (locus TcSNP139). Observed heterozygosity (Ho) showed a mean of 0.27 across loci, with individual values spanning 0.039–0.901, while expected heterozygosity (He) averaged 0.39 and ranged from 0.038 to 0.500. The minor-allele frequency (MAF) averaged 0.31 per marker, with the lowest MAF of 0.020 at locus TcSNP277 and the highest of 0.496 at locus TcSNP 878.

Genetic diversity in Panama was broadly comparable to that observed in the international reference collection (Table S3). Expected heterozygosity was similar between Panamanian cacao and reference accessions (He = 0.384 and 0.393, respectively), whereas observed heterozygosity was higher in Panama (Ho = 0.272 vs. 0.161). The inbreeding coefficient was lower in Panama (FIS = 0.281) than in the reference collection (FIS = 0.596), indicating lower levels of heterozygote deficit among Panamanian accessions.

### 3.3 Genetic ancestry and population structure of Panamanian cacao

The genetic relationships among the Panamanian *Theobroma cacao* samples and the reference accessions are presented in the principal coordinate analysis (PCoA) plot and the UPGMA dendrogram (Figure 3A–B). The first two PCoA axes explained 28.37% and 16.33% of the total genetic variation, respectively. Reference genotypes grouped into ten distinct clusters corresponding to known genetic groups: Criollo, Amelonado, Contamana, Curaray, Guiana, Iquitos, Marañón, Nanay, Nacional, and Purús.

**Figure 3.**
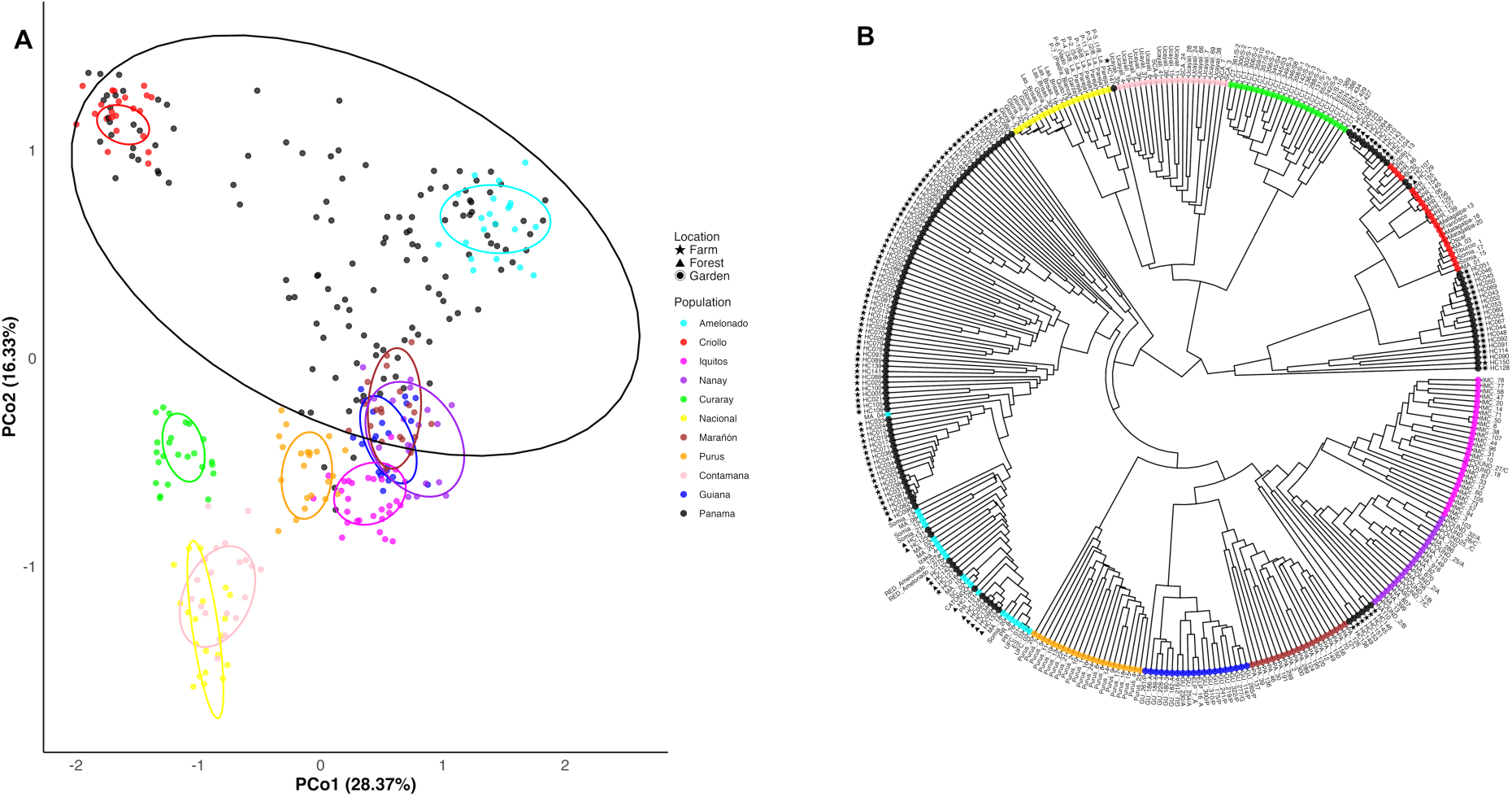
Genetic structure of Panamanian cacao and reference accessions. (A) Principal coordinate analysis (PCoA) showing genetic relationships among 128 Panamanian *Theobroma cacao* accessions and 219 reference genotypes. The first and second axes explain 28.37% and 16.33% of the total genetic variation, respectively. (B) UPGMA dendrogram based on SNP-derived genetic dissimilarities illustrating genetic relationships among Panamanian and reference accessions. Population assignments are color-coded consistently across both panels.

Several Panamanian samples overlapped with the Criollo and Amelonado clusters in the PCoA, while others were scattered near Upper Amazonian groups such as Iquitos, Nanay, and Contamana. A large number of Panamanian accessions occupied intermediate positions across the plot, suggesting varying levels of admixture and diverse ancestry. No Panamanian individuals were found near the Nacional or Guiana clusters, which remained genetically distinct from the Panamanian material.

The UPGMA dendrogram (Figure 3B) showed clustering patterns broadly consistent with those observed in the PCoA. One major cluster contained Nacional, Guiana, and Purús reference accessions, whereas Panamanian samples grouped primarily with Criollo, Amelonado, and several Upper Amazonian genetic groups. Within this cluster, several Panamanian accessions grouped closely with Criollo and Amelonado reference genotypes, while others occupied intermediate positions, reflecting varying degrees of admixture. Notably, the Panamanian accessions most closely associated with Criollo references were collected primarily from gardens and forests.

The symbols in the dendrogram further highlight the ecological context of collection. Samples from farms, forests, and home gardens were distributed throughout the dendrogram. Accessions from all three environments occurred within the Criollo and Amelonado-associated clusters; however, forest and garden accessions were more frequently associated with the Criollo cluster.

Bayesian clustering analysis using STRUCTURE at K = 10 clearly distinguished the same ten ancestral genetic groups among reference accessions (Fig. 4A), which served as the baseline for inferring population structure in Panamanian accessions.

**Figure 4.**
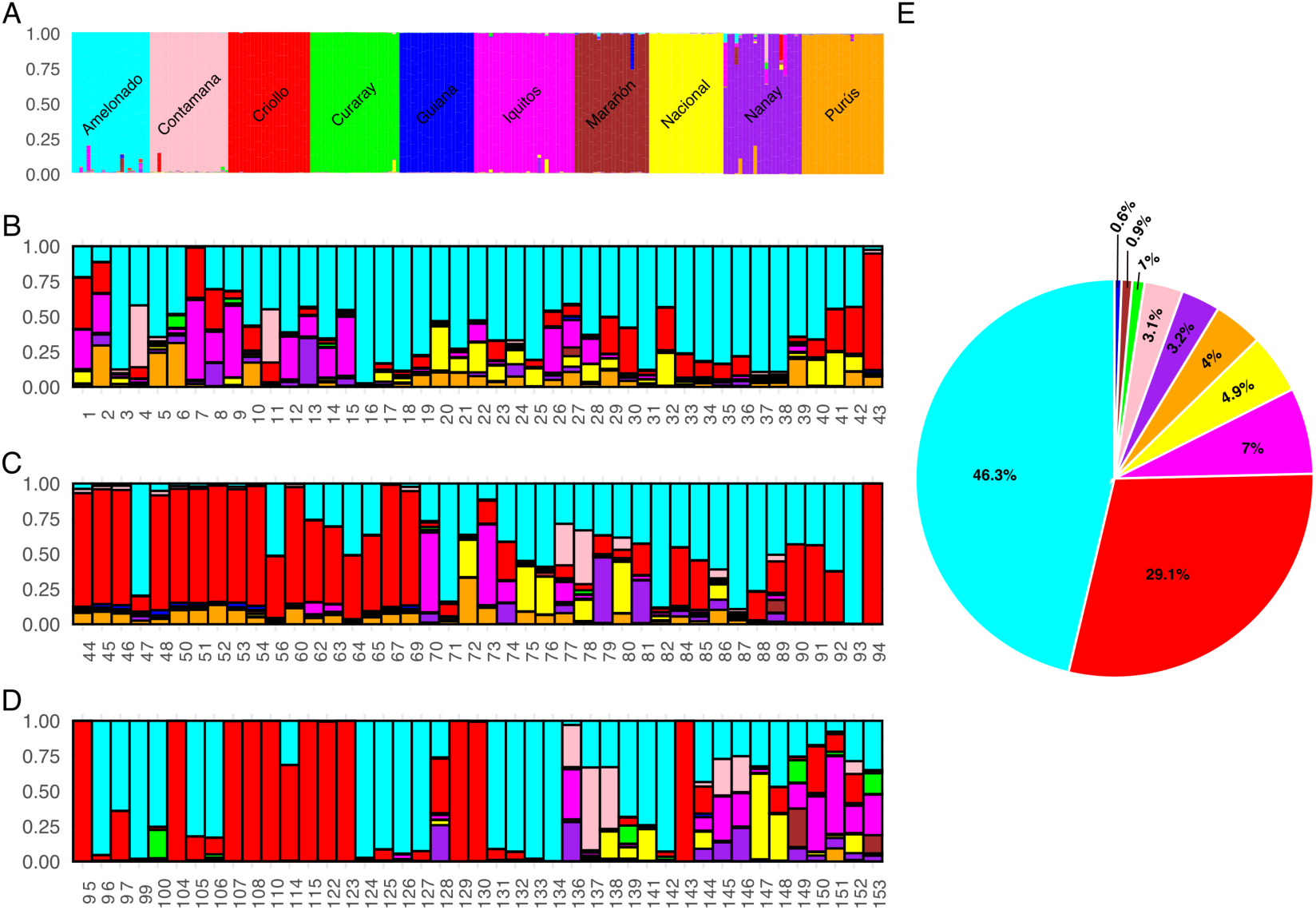
Genetic ancestry of Panamanian cacao: individual-level admixture and population-level ancestry proportions using STRUCTURE (K = 10). Each vertical bar represents an individual genotype, and the colors indicate estimated membership proportions to ten ancestral genetic groups. A) shows the reference populations. (B–D) display Panamanian accessions sorted by STRUCTURE-assigned group membership. E) summarizes average ancestry contributions of each genetic group across all Panamanian samples using a pie chart

Panamanian accessions were then sorted by STRUCTURE-inferred ancestry (Figs. 4B–D), revealing considerable variation in genetic background across sites. Genotypes were considered of single ancestry if one cluster’s membership proportion (Q) was ≥ 0.75 with no other group contributing more than Q = 0.25. By this criterion, 45.3% of Panamanian samples were classified as single ancestry genotypes, primarily from the Amelonado (25.8%) and Criollo (19.5%) groups. The remaining 54.7% of accessions were identified as admixed, reflecting hybrid ancestry involving two or more genetic groups. Among the admixed individuals, 7 accessions showed predominant admixture between Amelonado and Criollo ancestry, consistent with Trinitario-like ancestry. The remaining admixed individuals showed ancestry contributions from three or more genetic groups.

The summary of average ancestry proportions across all Panamanian accessions (Fig. 4E) confirmed the dominance of Amelonado ancestry (46.3%), followed by Criollo (29.1%), with minor contributions from Purús (7%), Nacional (5.6%), and Curaray (4.5%). These results highlight a complex ancestry structure within Panama, where traditional cacao lineages are preserved alongside high levels of admixture.

### 3.4 Human-managed agroecosystems harbor distinct cacao genetic resources

To evaluate whether ancestry composition differed among management systems, cacao accessions were classified according to their collection environment as farms, forests, or home gardens. Penalized multinomial logistic regression revealed clear structuring of ancestry composition across habitat types and geographic gradients, with Criollo ancestry enriched in home gardens and Amelonado ancestry predominating in farms (Figure 5). As described in Methods, Firth penalized logistic regression (adjusted for longitude) was used as the primary inferential model due to quasi-complete separation in the farm category. After adjustment, Criollo ancestry remained significantly associated with both home gardens (OR = 76.82, 95% CI [9.75, 9929.6]; p < 0.001) and forest habitats (OR = 20.68, 95% CI [1.76, 2934.2]; p = 0.013) relative to farms (Table S4). Comparison with the unadjusted model showed that the forest odds ratio decreased substantially after adjustment (66.7 to 20.7), indicating that part of the raw forest association reflected the underlying longitudinal gradient rather than habitat type alone; the garden association was comparatively robust to this adjustment (93.6 to 76.8; Table S4).

**Figure 5.**
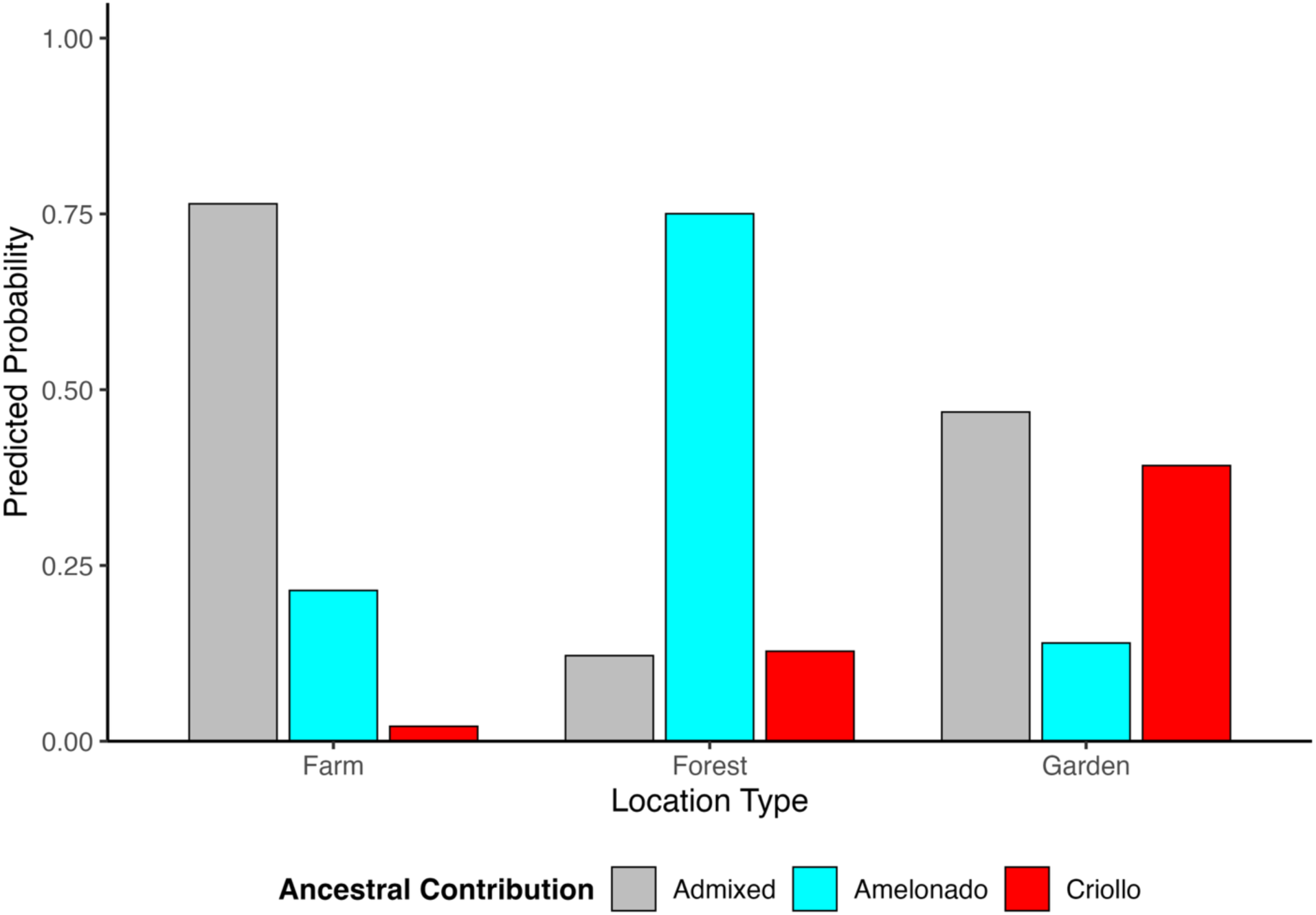
Habitat-specific predicted ancestry probabilities in Panamanian cacao. Predicted probabilities of *Theobroma cacao* accessions being assigned to Amelonado, admixed, or Criollo ancestry across farms, forests, and home gardens. Probabilities were estimated from a multinomial logistic regression model using STRUCTURE-based ancestry assignments and holding longitude at its mean value. Individuals with ancestry coefficients Q ≥ 0.75 for a single genetic group were classified as Amelonado or Criollo, whereas individuals below this threshold were classified as admixed.

A geographic trend was also detected along the longitudinal gradient. The predicted probability of Criollo ancestry increased eastward, particularly in garden habitats, where values exceeded 0.8 in the easternmost sampling locations. Conversely, the probability of Amelonado ancestry declined with increasing longitude (Figure S2; Table S6). Together, these results indicate that human-managed agroecosystems, especially home gardens, play a disproportionate role in maintaining Criollo-associated genetic resources within Panama.

### 3.5 Climate suitability and future conservation priorities

Climate suitability models showed very high discriminatory performance for both current and future scenarios (AUC > 0.99; Table S8). Among the retained bioclimatic predictors, isothermality (BIO3) and temperature seasonality (BIO4) were the most influential variables driving model predictions. Under current climatic conditions, BIO3 contributed most strongly to suitability estimates, whereas BIO4 became the dominant predictor in future projections. Because extremely high AUC values may indicate strong separation between presences and pseudo-absences or residual spatial bias, these results should be interpreted as evidence of model discrimination rather than definitive proof of ecological accuracy.

Under current climatic conditions, highly suitable areas for cacao cultivation were concentrated in Bocas del Toro, Colón, Panamá, and Herrera, with additional areas of moderate suitability in Veraguas, Ngäbe-Buglé, and Darién (Figure 6A). These regions largely coincide with areas where cacao is currently cultivated and where much of the genetic diversity identified in this study was sampled.

**Figure 6.**
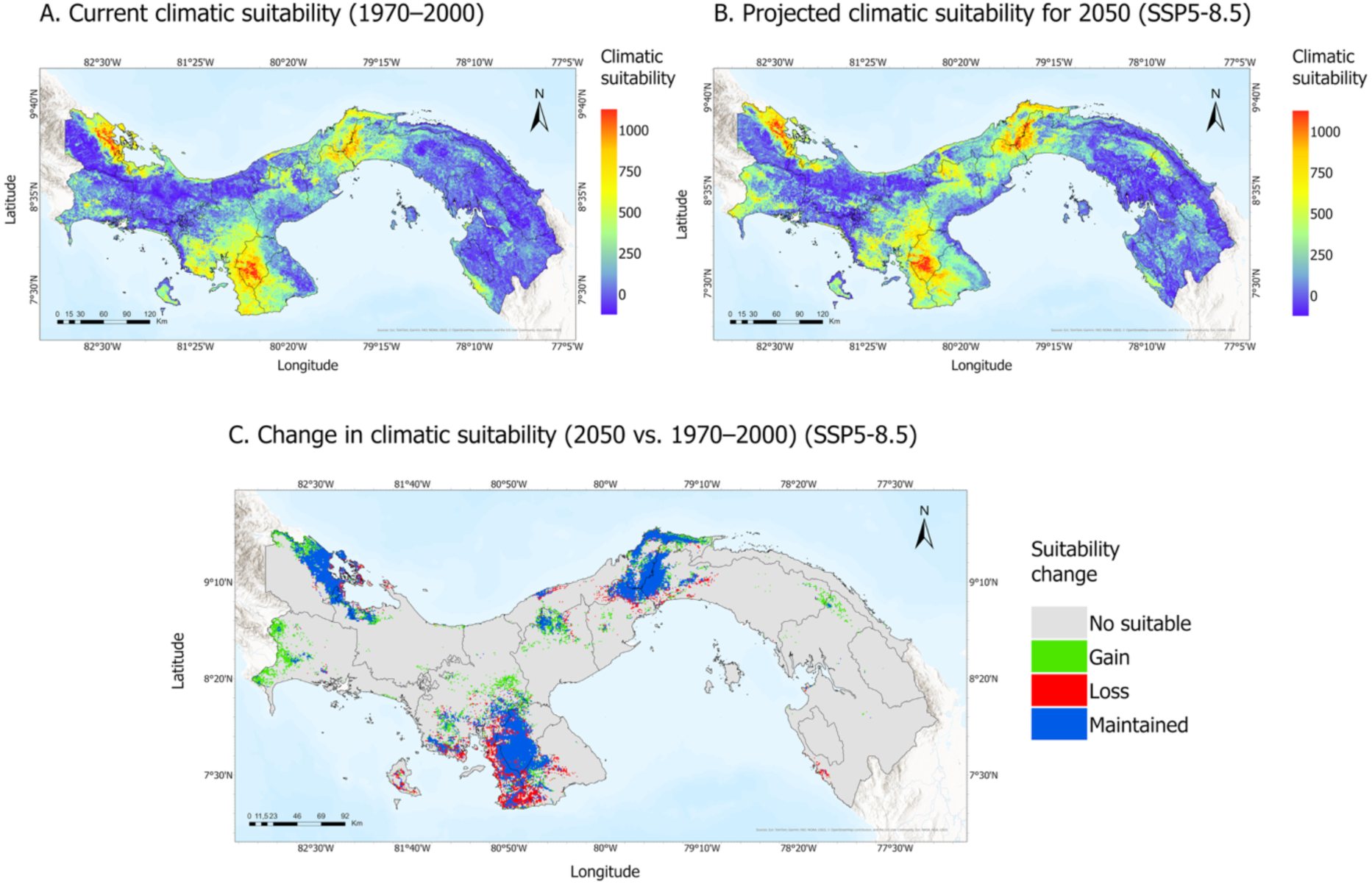
Climate suitability models for *Theobroma cacao* in Panama under current and future conditions. (A) Current climatic suitability (1970–2000), where the color gradient represents low (blue) to high (red) suitability for cacao cultivation. (B) Projected climatic suitability for 2050 under the high-emissions scenario SSP5-8.5. (C) Predicted changes in suitability between current and future conditions. Gray indicates areas unsuitable under both scenarios, green represents gains in suitability, red indicates losses, and blue denotes areas that remain suitable under both current and future climates.

Future projections under the SSP5-8.5 scenario indicated substantial spatial stability in the distribution of suitable habitat but also revealed important regional shifts (Figure 6B–C). Total climatically suitable area increased from 8,779.9 km² to 9,825.0 km², equivalent to an increase of approximately 11.9%. Suitability gains were concentrated in Chiriquí, Veraguas, and northwestern Darién, while losses were projected along the Pacific coast of Veraguas and the southwestern portion of Los Santos Province. Although localized reductions were also detected in Bocas del Toro and Panamá, these were less extensive than those observed in the central Pacific region.

Areas of future climatic suitability did not fully overlap with the current distribution of cacao genetic resources. Regions such as Bocas del Toro and Darién remain important reservoirs of cacao diversity, whereas Chiriquí and central Veraguas may become increasingly suitable for future cultivation. These patterns suggest that conservation planning should distinguish between areas that currently harbor valuable genetic resources and areas that may become climatically favorable for future production. Priority regions should therefore be identified by integrating genetic value, projected climatic suitability, sampling coverage, and feasibility of *in situ* or *ex situ* conservation.

## 4. Discussion

### 4.1 Human-managed agroecosystems maintain distinct cacao genetic resources in Panama

Our results indicate that cacao genetic diversity in Panama is associated with human-managed agroecosystems. The predominance of Amelonado and Criollo ancestries, together with a high proportion of admixed individuals, is consistent with centuries of introductions and farmer-mediated selection. Criollo genotypes were substantially more likely to occur in home gardens than in farms, even after accounting for the longitudinal ancestry gradient, suggesting that traditional management systems may maintain germplasm that is uncommon in commercial production systems. The wide profile-likelihood confidence intervals reflect sparse data and the complete absence of Criollo-assigned accessions in the farm category; effect magnitudes should therefore be interpreted with caution, particularly given the limited number of accessions available per habitat category.

In addition to this habitat structuring, Panamanian cacao exhibited moderate genetic diversity (He = 0.384), a positive inbreeding coefficient (FIS = 0.281), and a relatively high proportion of duplicate genotypes (20.9%), particularly in Herrera, where nearly half of sampled trees shared identical SNP profiles. This level of diversity is broadly comparable to farmer-maintained cacao populations elsewhere in Central America; Honduran and Nicaraguan accessions show lower observed heterozygosity (Ho = 0.206) but similar expected heterozygosity (He = 0.367; Ji et al., 2012). These patterns suggest genetic bottlenecks, clonal propagation, and restricted gene flow. Denser genomic approaches would help confirm whether accessions identified as genetically identical with the current 96-SNP panel represent true clonal duplicates or closely related individuals that this marker density cannot fully distinguish. Similar patterns have been reported in other cacao collections dominated by long-term vegetative propagation and limited germplasm exchange.

Traditional agroecosystems preserve locally adapted germplasm through continuous cultivation, and cultural selection rather than formal conservation programs (Galluzzi et al., 2010). The persistence of Criollo-dominated trees is particularly notable given their susceptibility (Cornejo et al., 2018). Under production-oriented management, such genotypes would be expected to decline, suggesting that factors beyond agronomic performance contribute to their persistence. The continued presence of Criollo ancestry in home gardens likely reflects cultural preferences and long-term management practices that have shaped cacao diversity across generations.

In contrast, Amelonado ancestry dominates farms. Amelonado germplasm became widely established throughout tropical America because of its adaptation to commercial production, illustrating how agricultural systems can selectively favor particular ancestry groups. Together, these findings show that farms and home gardens contribute complementary components of Panama’s cacao diversity. Commercial systems maintain productive lineages, whereas home gardens preserve culturally valued and potentially underrepresented genetic diversity.

### 4.2 Historical legacies continue to shape contemporary cacao populations

The genetic structure of Panamanian cacao reflects a long history of human-mediated dispersal, exchange, and cultivation. The predominance of admixed individuals indicates repeated episodes of genetic mixing rather than long-term isolation, consistent with growing evidence that cacao domestication involved multiple lineages and extensive exchanges across the Neotropics (Argout et al., 2023; Lanaud et al., 2024; Motamayor et al., 2008).

Archaeological and archaeogenomic evidence confirms the antiquity of cacao use in Panama (Section 3.1). More recently, archaeogenomic analyses revealed ancestry components related to modern Amelonado and Criollo groups, supporting the hypothesis that long-distance exchange networks facilitated genetic mixing among cacao populations thousands of years ago (Lanaud et al., 2024). These findings challenge the notion that ancient cacao cultivation was dominated by a single genetic lineage and instead point to a complex history of dispersal and admixture.

Panama’s position as an isthmian bridge likely facilitated these processes. Pre-Columbian societies maintained exchange networks linking present-day Panama with neighboring regions of Costa Rica and Colombia (Cooke & Sánchez, 2003), providing opportunities for the movement of crops and associated genetic diversity.

More recent historical processes further shaped contemporary populations. These well-documented colonial and commercial-era introductions (Section 3.1) likely reinforced existing admixture patterns while increasing the representation of particular ancestry groups within agricultural landscapes.

On Barro Colorado Island, Criollo- and Amelonado-associated trees occur within protected forest sampled in the Smithsonian’s 50-hectare plot. Archaeological evidence indicates prehistoric occupation of this area between approximately 1,700–1,300 and 700–400 years ago (Dent & Elsy, 2024), and since 1923 the island has remained protected, allowing relict cacao trees to persist within a regenerating forest matrix rather than as naturally occurring wild populations. Similar patterns in Darién suggest some forest cacao populations originated from historical agroforestry systems that persisted through ecological succession.

Together, these findings indicate that contemporary cacao populations are the cumulative outcome of ancient exchange networks, historical cultivation practices, and modern agricultural development. The genetic structure observed today therefore reflects a dynamic history of interactions between people and landscapes that has shaped cacao diversity across centuries.

### 4.3 Climate change is projected to redistribute suitable areas, but current genetic diversity hotspots are not all located within future refugia

Climate change is expected to reshape the distribution of climatically suitable areas for cacao in Panama, creating both opportunities and challenges for long-term conservation. Consistent with Results (Section 3.5), these shifts were spatially heterogeneous rather than uniform, in line with broader regional assessments indicating that climate change is likely to redistribute rather than uniformly reduce cacao-growing areas (Bunn et al., 2019; Schroth et al., 2016). In contrast, most comparable assessments report that projected gains in cacao suitability are typically modest or substantially reduced once land-use constraints are applied, or find net declines under climate change (Ariza-Salamanca et al., 2023; Ceccarelli et al., 2021; Gloy et al., 2025).The larger net increase projected here for Panama should therefore be interpreted with some caution, and may partly reflect the finer spatial resolution and national scale of our analysis.

Projected losses in the Azuero Peninsula align with intensification of the Panama Low-Level Jet and ITCZ-driven atmospheric subsidence reducing Pacific-slope rainfall (Ordonez-Zuniga et al., 2021), whereas gains in northwestern Chiriquí and east-central Veraguas reflect the orographic rain shadow of the Central Cordillera, where future rainfall reductions could favor more defined microseasonality suitable for cacao (Fábrega et al., 2013; World Bank Group, 2024). Provinces retaining high suitability, including Bocas del Toro, Colón, and Panamá, are instead shaped by Hadley Cell circulation and Caribbean trade winds, which maintain comparatively low rainfall seasonality (Taylor et al., 2012).

These climatic suitability models were based primarily on bioclimatic variables and should therefore be interpreted as estimates of climatic suitability rather than direct predictions of future cultivation; they did not explicitly incorporate soil properties, pest and disease pressure, water availability, current forest cover, protected-area status, or farmer decision-making, all of which can strongly influence the realized distribution of cacao production systems. Model outputs are also sensitive to the occurrence records used, the spatial distribution of sampling effort, predictor collinearity filtering, pseudo-absence design, the calibration algorithm, and the choice of climate scenario and general circulation models. Moreover, high discrimination metrics such as AUC (Table S8) do not necessarily guarantee ecological accuracy when presence records are spatially clustered. Projected gains or losses under SSP5-8.5 should therefore be interpreted cautiously. Importantly, future climatic suitability does not necessarily coincide with the current distribution of genetic diversity. Some components of Panama’s cacao diversity occur in home gardens, remnant forest patches, and historically managed landscapes that may fall outside future areas of optimal suitability. These genetic resources could therefore become increasingly vulnerable even if the total area suitable for cultivation remains stable or expands. Conservation will require more than maintaining suitable environments for production; it will also mean protecting the specific populations and lineages that harbor valuable genetic variation.

The projected persistence of suitable conditions in parts of Bocas del Toro is particularly noteworthy because this region currently supports populations representing multiple ancestry groups. Consequently, Bocas del Toro may remain important not only for future cacao production (Bunn et al., 2019), but also for the conservation of genetic diversity under changing climatic conditions, given its role as a documented reservoir of admixed and Criollo-associated ancestry. At the same time, newly suitable areas may create opportunities for cultivation in regions that currently play a limited role in national production. However, adaptation strategies based solely on geographic relocation could inadvertently contribute to genetic homogenization if new plantations rely on a limited number of commercial cultivars. These models do not account for local adaptation, phenotypic plasticity, or adaptive management by farmers, all of which may buffer some populations against projected climatic change. Integrating climate projections with genetic information will therefore remain essential for identifying priority areas where both cacao cultivation and its underlying genetic resources are most likely to persist.

### 4.4 Implications for conservation and climate-resilient cacao systems: *integrating in situ* and *ex situ* strategies

Given the historical, genetic, and climatic patterns identified here, conservation strategies for cacao in Panama should integrate the preservation of germplasm with the protection of the agroecosystems and management practices through which diversity has been maintained.

An important observation was that none of the Panamanian accessions assigned to the Criollo genetic group showed an identical 96-SNP profile to any of the reference Criollo genotypes included in this study, including Criollo 13. This accession, maintained at CATIE in Costa Rica, is documented as originating from Panama in the institution’s germplasm catalogue (Morera et al., 1991) and has been widely used as a representative of the Criollo genetic group in population genetic and genomic studies (Cosme et al., 2016; Lindo et al., 2018; Motamayor et al., 2008; B. Wang et al., 2020). Hierarchical clustering further showed that a subset of Panamanian accessions grouped within the same clade as the reference Criollo genotypes despite lacking an identical SNP profile, whereas the majority of Panamanian Criollo-assigned accessions clustered on a separate branch that did not include any reference genotype (Figure 3). This heterogeneous clustering pattern, together with the absence of identical multilocus matches, suggests that the Criollo-associated germplasm currently sampled in Panama differs, at least in part, from the reference accessions included in this study. Broader genomic analyses will be required to determine whether this pattern reflects previously undocumented regional variation within the Criollo genetic group or finer-scale differentiation within existing Criollo diversity.

The strong association between Criollo ancestry and home gardens further emphasizes the conservation value of traditional agroecosystems. These systems maintain culturally important cacao varieties and genetic combinations accumulated through centuries of local selection, exchange, and cultivation. As a result, these systems conserve genetic diversity alongside the social and cultural practices that have sustained it.

The absence of a national cacao germplasm collection in Panama represents a significant conservation gap. While international collections such as CATIE and USDA-TARS play critical roles in preserving global cacao diversity, they may not adequately capture locally differentiated populations maintained within Panamanian landscapes.

Based on our findings, three conservation priorities emerge for Panama. First, a national cacao germplasm bank should be established to preserve genetic resources currently maintained in farms, forests, and home gardens. Second, traditional home gardens should be formally incorporated into national cacao conservation strategies because they harbor a disproportionate fraction of Criollo-associated diversity. Third, conservation planning should prioritize regions such as Bocas del Toro and Darién, which combine high genetic value with projected climatic suitability, while also promoting the collection of underrepresented Criollo-associated germplasm from traditional agroecosystems. Together, these actions would strengthen the long-term conservation of Panamanian cacao genetic resources while supporting adaptation to future climatic conditions.

These priorities align with Panama’s National Cacao Law (Law 261, 2021) and its implementing decree, which created the Oficina Nacional del Cacao to coordinate technical assistance and sector-wide monitoring. The genetic and climatic evidence presented here could inform this office’s prioritization criteria across all cacao-growing provinces, identifying regions and management systems that merit early germplasm collection based on combined genetic value and climatic stability, rather than production volume alone.

Maintaining traditional production systems, Indigenous cultivation practices, and smallholder-managed gardens is therefore as important as preserving seeds or clones in repositories. Integrating in situ and ex situ conservation approaches will protect existing diversity while sustaining the evolutionary and cultural processes that contribute to its persistence.

### 4.5 Conclusions

Cacao genetic resources in Panama are strongly linked to human-managed agroecosystems. Accessions with predominant Criollo ancestry were markedly more likely to occur in home gardens and forest remnants than on farms, even after accounting for longitudinal variation in ancestry, whereas farm accessions were dominated by Amelonado ancestry. Traditional home gardens and historically managed landscapes are therefore not merely sites of production but important reservoirs of genetic diversity shaped by generations of cultivation and management. Contemporary genetic patterns reflect the cumulative influence of ancient dispersal, colonial persistence, and modern agricultural development. Notably, Panamanian Criollo-assigned accessions did not show identical 96-SNP profiles to the international reference genotypes examined, suggesting that regional germplasm surveys can reveal locally differentiated diversity that may be underrepresented in existing collections. Climate change is projected to redistribute suitable cultivation areas, with expansion in Chiriquí and Veraguas and losses in the Azuero Peninsula, but future climatic suitability does not necessarily coincide with current genetic diversity hotspots. Climate-resilient cacao conservation in Panama will therefore require integrating genetic, historical, and climatic evidence to protect germplasm together with the agroecosystems and cultural practices that sustain it.

## Acknowledgments

This work was supported by the Secretaría Nacional de Ciencia, Tecnología e Innovación de Panamá (SENACYT; Grant FID17-104 and SENACYT Ph.D. Scholarship to H.E.C), with additional support from a donation by Darlene Bookhoff, and L.C.M. acknowledges support from the SENACYT SNI program. The authors thank IDIAP, Smithsonian ForestGEO, MiAMBIENTE, and Reserva Natural Punta Patiño for facilitating access to sample collection sites. We are especially grateful to Dr. Dapeng Zhang for his support with SNP genotyping and for providing access to reference genotype datasets used in this study. We sincerely thank the local communities and cacao growers who generously shared their knowledge and facilitated access to sampling sites.

We also thank Yaleskia Valdés, Charles Martin, Eliab Ramos, Domingo Diaz, Abiel Gutierrez and Pascal Artieda for their invaluable assistance during field surveys and sample collection.

## Author Contributions

H.E.C. and L.C.M. conceived and coordinated the project. H.E.C. and L.C.M. conducted field sampling, H.E.C. analyzed the SNP data, interpreted the genetic results, prepared the figures, contributed to the historical review and interpretation of historical sources, and wrote the original manuscript draft. D.M.J. contributed to climate suitability modeling and interpretation of environmental data. All authors reviewed, edited, and approved the final manuscript.

## Data Availability Statement

The data supporting the findings of this study are available within the article and its Supporting Information files. Additional data are available from the corresponding author.

## References

Albert, A., & Anderson, J. A. (1984). On the existence of maximum likelihood estimates in logistic regression models. Biometrika, 71(1), 1–10. 10.1093/biomet/71.1.1

Arel-Bundock, V., Enevoldsen, N., & Yetman, C. (2018). countrycode: An R package to convert country names and country codes. Journal of Open Source Software, 3(28), 848. 10.21105/joss.00848

Arevalo-Gardini, E., Meinhardt, L. W., Zuñiga, L. C., Arévalo-Gardni, J., Motilal, L., & Zhang, D. (2019). Genetic identity and origin of “Piura Porcelana”—A fine-flavored traditional variety of cacao (Theoborma cacao) from the Peruvian Amazon. Tree Genetics & Genomes, 15(1), 11. 10.1007/s11295-019-1316-y

Argout, X., Droc, G., Fouet, O., Rouard, M., Labadie, K., Rhoné, B., Loor, G. R., & Lanaud, C. (2023). Pangenomic exploration of Theobroma cacao: New Insights into Gene Content Diversity and Selection During Domestication. bioRxiv. 10.1101/2023.11.03.565324

Ariza-Salamanca, A. J., Navarro-Cerrillo, R. M., Quero-Pérez, J. L., Gallardo-Armas, B., Crozier, J., Stirling, C., de Sousa, K., & González-Moreno, P. (2023). Vulnerability of cocoa-based agroforestry systems to climate change in West Africa. Scientific Reports, 13, 10033. 10.1038/s41598-023-37180-3

Barbet-Massin, M., Jiguet, F., Albert, C. H., & Thuiller, W. (2012). Selecting pseudo-absences for species distribution models: How, where and how many? Methods in Ecology and Evolution, 3(2), 327–338. 10.1111/j.2041-210X.2011.00172.x

Bartley, B. G. D. (2005). The Genetic Diversity of Cacao and Its Utilization. CABI Publishing.

Bennett, H. H., & Taylor, W. A. (1912). The agricultural possibilities of the Canal Zone: Pt. I. Reconnaissance soil survey. U.S. Department of Agriculture.

Bourgois, P. I. (1994). Banano, etnia y lucha social en Centro América. Editorial Departamento Ecuménico de Investigaciones.

Bunn, C., Fernández-Kolb, P., Wiegel, J., Guharay, F., Hurtado, N., Castro-Llanos, F., & Lundy, M. (2019). Climate Change Impacts on Cocoa Production in Central America and the Caribbean. International Center for Tropical Agriculture (CIAT). 10.7910/DVN/QUKZTO

Castillero Calvo, A. (2017). Conquista, evangelización y resistencia (2nd ed.). Editora Novo Art.

Castillero Calvo, A. (2019). Panamá: Síntesis de su historia colonial (2nd, Ed.). Fundación Biblioteca Ayacucho.

Ceccarelli, V., Fremout, T., Zavaleta, D., Lastra, S., Imán Correa, S., Arévalo-Gardini, E., Rodriguez, C. A., Cruz Hilacondo, W., & Thomas, E. (2021). Climate change impact on cultivated and wild cacao in Peru and the search of climate change-tolerant genotypes. Diversity and Distributions, 27(8), 1462–1476. 10.1111/ddi.13294

Ceccarelli, V., Lastra, S., Loor Solórzano, R. G., Chacón, W. W., Nolasco, M., Sotomayor Cantos, I. A., Plaza Avellán, L. F., López, D. A., Fernández Anchundia, F. M., Dessauw, D., Orozco-Aguilar, L., & Thomas, E. (2022). Conservation and use of genetic resources of cacao (Theobroma cacao L.) by gene banks and nurseries in six Latin American countries. Genetic Resources and Crop Evolution, 69(3), 1283–1302. 10.1007/s10722-021-01304-3

Centro Agronómico Tropical de Investigación y Enseñanza (CATIE). (2018). El sector cacao en Centroamérica: Estado de desarrollo en el año 2007 – Línea de base del Proyecto Cacao Centroamérica (PCC). CATIE.

Cerda, R., & Orozco, S. (2009). Foro: Genética del cacao en Panamá: ¿Qué estamos haciendo y hacia dónde vamos? Universidad Tecnológica de Panamá. https://www.yumpu.com/es/document/view/30206334/foro-genactica-del-cacao-en-panama-catie

Clopper, C. J., & Pearson, E. S. (1934). The use of confidence or fiducial limits illustrated in the case of the binomial. Biometrika, 26(4), 404–413. 10.1093/biomet/26.4.40

COCABO. (2025). Historia de la cooperativa. Cooperativa de Servicios Múltiples Cacao Bocatoreña, R.L. https://cocabo.org/

Coe, S. D., & Coe, M. D. (1996). The True History of Chocolate. Thames and Hudson

Cooke, R. G., & Sánchez, L. (2003). Panamá precolombino. In F. W. Lange (Ed.), Historia de Panamá (pp. 23–71). Presidencia de la República.

Cornejo, O. E., Yee, M.-C., Dominguez, V., Andrews, M., Sockell, A., Strandberg, E., Livingstone, D., Stack, C., Romero, A., Umaharan, P., Royaert, S., Tawari, N. R., Ng, P., Gutierrez, O., Phillips, W., Mockaitis, K., Bustamante, C. D., & Motamayor, J. C. (2018). Population genomic analyses of the chocolate tree, Theobroma cacao L., provide insights into its domestication process. Communications Biology, 1(1), 167. 10.1038/s42003-018-0168-6

Cosme, S., Cuevas, H. E., Zhang, D., Oleksyk, T. K., & Irish, B. M. (2016). Genetic diversity of naturalized cacao (Theobroma cacao L.) in Puerto Rico. Tree Genetics & Genomes, 12(5), 88. 10.1007/s11295-016-1045-4

Dent, D. H., & Elsy, A. (2024). Structure, Diversity, and Composition of Secondary Forests of the Barro Colorado Nature Monument. Smithsonian Institution Scholarly Press. 10.5479/si.26809615

Dickau, R. (2010). Microbotanical and Macrobotanical Evidence of Plant Use and the Transition to Agriculture in Panama. In A. M. VanDerwarker & T. M. Peres (Eds.), Integrating Zooarchaeology and Paleoethnobotany: A Consideration of Issues, Methods, and Cases (pp. 99–134). Springer. 10.1007/978-1-4419-0935-0_6

Fábrega, J., Nakaegawa, T., Pinzón, R., Nakayama, K., Arakawa, O., & SOUSEI Theme-C Modeling Group. (2013). Hydroclimate projections for Panama in the late 21st Century. Hydrological Research Letters, 7(2), 23–29. 10.3178/hrl.7.23

Fernández de Oviedo, G. (1851). Historia general y natural de las Indias (Vol. 1). Real Academia de la Historia.

Ferro-Lozano, N., Mejia, L. C., Kuyper, T. W., & Bonnema, G. (2026). Contrasting germplasm composition and propagation practices in the two major cacao-growing areas in Panama. Genetic Resources and Crop Evolution, 73, 234. 10.1007/s10722-026-02844-2

Fick, S. E., & Hijmans, R. J. (2017). WorldClim 2: New 1-km spatial resolution climate surfaces for global land areas. International Journal of Climatology, 37(12), 4302–4315. 10.1002/joc.5086

Firth, D. (1993). Bias reduction of maximum likelihood estimates. Biometrika, 80(1), 27–38. 10.1093/biomet/80.1.27

Friedman, J., Hastie, T., & Tibshirani, R. (2010). Regularization Paths for Generalized Linear Models via Coordinate Descent. Journal of Statistical Software, 33(1), 1–22. 10.18637/jss.v033.i01

Galluzzi, G., Eyzaguirre, P., & Negri, V. (2010). Home gardens: Neglected hotspots of agro-biodiversity and cultural diversity. Biodiversity and Conservation, 19(13), 3635–3654. 10.1007/s10531-010-9919-5

Gloy, N., Chemura, A., Tchindjang, M., & Gornott, C. (2025). Mapping climate change impacts on cocoa agroforestry systems in Cameroon to mitigate future deforestation. Agronomy for Sustainable Development, 45, 66. 10.1007/s13593-025-01042-1

Gobierno de Panamá. (2022). Decreto Ejecutivo No. 15 de 14 de julio de 2022 que reglamenta la Ley 261 de 23 de diciembre de 2021 (Issues 29612-B).

Gutiérrez, A. (2020). Caracterización morfológica de tres genotipos criollos promisorios de Theobroma cacao L. en Panamá. Ciencia Agropecuaria, 30, 150–169.

Heinze, G., & Schemper, M. (2002). A solution to the problem of separation in logistic regression. Statistics in Medicine, 21(16), 2409–2419. 10.1002/sim.1047

Henderson, J. S., Joyce, R. A., Hall, G. R., Hurst, W. J., & McGovern, P. E. (2007). Chemical and archaeological evidence for the earliest cacao beverages. Proceedings of the National Academy of Sciences, 104(48), 18937–18940. 10.1073/pnas.0708815104

Hijmans, R., Barbosa, M., Bivand, R., Brown, A., Chirico, M., Cordano, E., Krzystof, D., Pebesma, E., Rowlingson, B., & Sumner, M. (2020). terra: Spatial Data Analysis [Computer software]. CRAN: Contributed Packages. 10.32614/CRAN.PACKAGE.TERRA

Hijmans, R., Barbosa, M., Ghosh, A., & Mandel, A. (2021). *geodata: Download Geographic Data* [Computer software]. CRAN: Contributed Packages. 10.32614/CRAN.PACKAGE.GEODATA

Huang, D., An, Q., Huang, S., Tan, G., Quan, H., Chen, Y., Zhou, J., & Liao, H. (2023). Biomod2 modeling for predicting the potential ecological distribution of three Fritillaria species under climate change. Scientific Reports, 13(1), 18801. 10.1038/s41598-023-45887-6

Ji, K., Zhang, D., Motilal, L. A., Boccara, M., Lachenaud, P., & Meinhardt, L. W. (2012). Genetic diversity and parentage in farmer varieties of cacao (Theobroma cacao L.) from Honduras and Nicaragua as revealed by single nucleotide polymorphism (SNP) markers. Genetic Resources and Crop Evolution, 60(2), 441–453. 10.1007/s10722-012-9847-1

Kalinowski, S. T., Manlove, K. R., & Taper, M. L. (2007). ONCOR: A computer program for genetic stock identification. Montana State University. https://www.montana.edu/kalinowski/software/ONCOR.htm

Kamvar, Z. N., Tabima, J. F., & Grünwald, N. J. (2014). poppr: An R package for genetic analysis of populations with clonal, partially clonal, and/or sexual reproduction. PeerJ, 2, e281. 10.7717/peerj.281

Lanaud, C., Vignes, H., Utge, J., Valette, G., Rhoné, B., Garcia Caputi, M., Angarita Nieto, N. S., Fouet, O., Gaikwad, N., Zarrillo, S., Powis, T. G., Cyphers, A., Valdez, F., Olivera Nunez, S. Q., Speller, C., Blake, M., Valdez, F. J., Raymond, S., Rowe, S. M., … Argout, X. (2024). A revisited history of cacao domestication in pre-Columbian times revealed by archaeogenomic approaches. Scientific Reports, 14(1), 2972. 10.1038/s41598-024-53010-6

Lindo, A. A., Robinson, D. E., Tennant, P. F., Meinhardt, L. W., & Zhang, D. (2018). Molecular Characterization of Cacao (Theobroma cacao) Germplasm from Jamaica Using Single Nucleotide Polymorphism (SNP) Markers. Tropical Plant Biology, 11(3–4), 93–106. 10.1007/s12042-018-9203-5

Lipper, L., Thornton, P., Campbell, B. M., Baedeker, T., Braimoh, A., Bwalya, M., Caron, P., Cattaneo, A., Garrity, D., Henry, K., Hottle, R., Jackson, L., Jarvis, A., Kossam, F., Mann, W., McCarthy, N., Meybeck, A., Neufeldt, H., Remington, T., & Torquebiau, E. F. (2014). Climate-smart agriculture for food security. Nature Climate Change, 4(12), 1068–1072. 10.1038/nclimate2437

Mata-Quirós, A., Arciniegas-Leal, A., Phillips-Mora, W., Meinhardt, L. W., Motilal, L., Mischke, S., & Zhang, D. (2018). Assessing hidden parentage and genetic integrity of the “United Fruit Clones” of cacao (Theobroma cacao) from Costa Rica using SNP markers. Breeding Science, 68(5), 545–553. 10.1270/jsbbs.18057

Morera, J., Paredes, A., & Mora, A. (1991). Germoplasma de cacao en el CATIE entre 1947 y 1991 (Nos. 0534–5391; Publicación Miscelánea A1/SC, pp. 1–49). IICA/CATIE. https://repositorio.iica.int/items/dc44ead5-cb32-412e-b0d5-b10737e92091

Motamayor, J. C., Lachenaud, P., Mota, J. W. da S. e, Loor, R., Kuhn, D. N., Brown, J. S., & Schnell, R. J. (2008). Geographic and Genetic Population Differentiation of the Amazonian Chocolate Tree (Theobroma cacao L). PLOS ONE, 3(10), e3311. 10.1371/journal.pone.0003311

Naimi, B. (2012). *usdm: Uncertainty Analysis for Species Distribution Models* [Computer software]. CRAN: Contributed Packages. 10.32614/CRAN.PACKAGE.USDM

Nieves-Orduña, H. E., Krutovsky, K. V., & Gailing, O. (2023). Geographic distribution, conservation, and genomic resources of cacao Theobroma cacao L. Crop Science, 63(4), 1750–1778. 10.1002/csc2.20959

Ordonez-Zuniga, S. A., Correa-Ramirez, M. A., Ricaurte-Villota, C., & Bastidas-Salamanca, M. (2021). The Panama Low Level Jet: Extension, annual cycle and modes of variation. Latin American Journal of Aquatic Research, 49(5), 750–762. 10.3856/vol49-issue5-fulltext-2591

Osorto Nuñez, M. H. (2022). Los modelos de cambio climático futuro como predictores de la reducción del área de distribución de dos especies de cíclidos endémicos de Honduras. Biología Acuática, 38, e028. 10.24215/16684869e028

Peakall, R., & Smouse, P. E. (2012). GenAlEx 6.5: Genetic analysis in Excel. Population genetic software for teaching and research—An update. Bioinformatics, 28(19), 2537–2539. 10.1093/bioinformatics/bts460

Pearson, K. G., & Holden, A. L. (1945). Manila (Abaca) Fiber for World War II. Proceedings, 71(5), 507.

Peterson, A. T., Papeş, M., & Soberón, J. (2008). Rethinking receiver operating characteristic analysis applications in ecological niche modeling. Ecological Modelling, 213(1), 63–72. 10.1016/j.ecolmodel.2007.11.008

Phillips-Mora, W. (2003). Origin, biogeography, genetic diversity and taxonomic affinities of the cacao (Theobroma cacao L.) fungus Moniliophthora roreri (Cif.) Evans et al. As determined using molecular, phytopathological and morpho-physiological evidence. University of Reading.

Phillips-Mora, W., Ortiz, C. F., & Aime, M. C. (2007). Fifty years of frosty pod rot in Central America: Chronology of its spread and impact from Panama to Mexico. I, 1039–1047.

Powis, T. G., Cyphers, A., Gaikwad, N. W., Grivetti, L., & Cheong, K. (2002). Cacao use and the San Lorenzo Olmec. Proceedings of the National Academy of Sciences, 99(25), 13395–13400. 10.1073/pnas.172370699

Pritchard, J. K., Stephens, M., & Donnelly, P. (2000). Inference of population structure using multilocus genotype data. Genetics, 155(2), 945–959. 10.1093/genetics/155.2.945

Reverte Coma, J. (1995). Indios, teribes y otras gentes de Panamá. Ministerio de Cultura.

Schliep, K. P. (2011). phangorn: Phylogenetic analysis in R. Bioinformatics, 27(4), 592–593. 10.1093/bioinformatics/btq706

Schroth, G., Läderach, P., Martinez-Valle, A. I., Bunn, C., & Jassogne, L. (2016). Vulnerability to climate change of cocoa in West Africa: Patterns, opportunities and limits to adaptation. Science of The Total Environment, 556, 231–241. 10.1016/j.scitotenv.2016.03.024

Sillero, N., Arenas-Castro, S., Enriquez-Urzelai, U., Vale, C. G., Sousa-Guedes, D., Martínez-Freiría, F., Real, R., & Barbosa, A. M. (2021). Want to model a species niche? A step-by-step guideline on correlative ecological niche modelling. Ecological Modelling, 456, 109671. 10.1016/j.ecolmodel.2021.109671

Sunny, A., Marmolejo, C., Vidal-López, R., Falconi-Briones, F. A., Cuervo-Robayo, Á. P., & Bolom-Huet, R. (2025). EcoNicheS: enhancing ecological niche modeling, niche overlap and connectivity analysis using the shiny dashboard and R package. PeerJ, 13, 1–29. 10.7717/peerj.23965

Taylor, G. T., Muller-Karger, F. E., Thunell, R. C., Scranton, M. I., Astor, Y., Varela, R., Ghinaglia, L. T., Lorenzoni, L., Fanning, K. A., & Hameed, S. (2012). Ecosystem responses in the southern Caribbean Sea to global climate change. Proceedings of the National Academy of Sciences, 109(47), 19315–19320. 10.1073/pnas.1207514109

The Canal Record. (1914, August 26). Official publication of the Panama Canal. The Canal Record, VIII(1). https://www.archive.org/details/panamacanalr8191415isth

Thuiller, W., Georges, D., Gueguen, M., Engler, R., Breiner, F., Lafourcade, B., Patin, R., & Blancheteau, H. (2025). *biomod2: Ensemble Platform for Species Distribution Modeling* [Computer software]. CRAN: Contributed Packages. 10.32614/CRAN.PACKAGE.BIOMOD2

Wang, B., Motilal, L. A., Meinhardt, L. W., Yin, J., & Zhang, D. (2020). Molecular Characterization of a Cacao Germplasm Collection Maintained in Yunnan, China Using Single Nucleotide Polymorphism (SNP) Markers. Tropical Plant Biology, 13(4), 359–370. 10.1007/s12042-020-09267-y

Wang, J., Singer, G., & Anderson, J. (2009). High-throughput SNP genotyping with the Fluidigm dynamic array system. Proceedings of the International Plant & Animal Genome XVII Conference.

Wickham, H. (2007). Reshaping data with the reshape package. Journal of Statistical Software, 21(12), 1–20. 10.18637/jss.v021.i12

Wickham, H., Chang, W., Henry, L., Pedersen, T. L., Takahashi, K., Wilke, C., Woo, K., Yutani, H., Dunnington, D., & van den Brand, T. (2025). *ggplot2: Create Elegant Data Visualisations Using the Grammar of Graphics* [Computer software]. CRAN: Contributed Packages. 10.32614/CRAN.PACKAGE.GGPLOT2

World Bank Group. (2024). Climate Risk Country Profile: Panama. World Bank Group. https://www.worldbank.org

Yu, G., Smith, D. K., Zhu, H., Guan, Y., & Lam, T. T. Y. (2017). ggtree: An R package for visualization and annotation of phylogenetic trees with their covariates and other associated data. Methods in Ecology and Evolution, 8(1), 28–36. 10.1111/2041-210X.12628

Zarrillo, S., Gaikwad, N., Lanaud, C., Powis, T. G., Viot, C., Lesur, I., & Blake, M. (2018). The use and domestication of Theobroma cacao during the mid-Holocene in the upper Amazon. Nature Ecology & Evolution, 2, 1879–1888. 10.1038/s41559-018-0697-x

Zhang, D., Martínez, W. J., Johnson, E. S., Somarriba, E., Phillips-Mora, W., Astorga, C., Mischke, S., & Meinhardt, L. W. (2012). Genetic diversity and spatial structure in a new distinct Theobroma cacao L. population in Bolivia. Genetic Resources and Crop Evolution, 59(2), 239–252. 10.1007/s10722-011-9680-y

Zhao, Z., Xiao, N., Shen, M., & Li, J. (2022). Comparison between optimized MaxEnt and random forest modeling in predicting potential distribution: A case study with Quasipaa boulengeri in China. Science of The Total Environment, 842, 156867. 10.1016/j.scitotenv.2022.156867

